# Hippocampal subfield vulnerability to α-synuclein pathology precedes neurodegeneration and cognitive dysfunction

**DOI:** 10.1101/2023.04.12.536572

**Authors:** Dylan J. Dues, An Phu Tran Nguyen, Katelyn Becker, Jiyan Ma, Darren J. Moore

**Affiliations:** Department of Neurodegenerative Science, Van Andel Institute, Grand Rapids, Michigan, USA; Chinese Institute for Brain Research, Beijing, China

**Author notes:** Corresponding author: Address: Van Andel Institute, <u>333 Bostwick Ave</u> NE, Grand Rapids, MI 49503, USA.

**Keywords:** Parkinson’s disease, Dementia with Lewy Bodies, Lewy pathology, alpha-synuclein, protein aggregation, dementia, neurodegeneration, animal models

## Abstract

Cognitive dysfunction is a salient feature of Parkinson’s disease (PD) and Dementia with Lewy bodies (DLB). The onset of dementia reflects the spread of Lewy pathology throughout forebrain structures. The mere presence of Lewy pathology, however, provides limited indication of cognitive status. Thus, it remains unclear whether Lewy pathology is the de facto substrate driving cognitive dysfunction in PD and DLB. Through application of α-synuclein fibrils *in vivo*, we sought to examine the influence of pathologic inclusions on cognition. Following stereotactic injection of α-synuclein fibrils within the mouse forebrain, we measured the burden of α-synuclein pathology at 1-, 3-, and 6-months post-injection within subregions of the hippocampus and cortex. Under this paradigm, the hippocampal CA2/3 subfield was especially susceptible to α- synuclein pathology. Strikingly, we observed a drastic reduction of pathology in the CA2/3 subfield across time-points, consistent with the consolidation of α-synuclein pathology into dense somatic inclusions followed by neurodegeneration. Silver-positive degenerating neurites were observed prior to neuronal loss, suggesting that this might be an early feature of fibril-induced neurotoxicity and a precursor to neurodegeneration. Critically, mice injected with α-synuclein fibrils developed progressive deficits in spatial learning and memory. These findings support that the formation of α-synuclein inclusions in the mouse forebrain precipitate neurodegenerative changes that recapitulate features of Lewy-related cognitive dysfunction.

**Highlights:**

- Mice injected with α-synuclein fibrils develop hippocampal and cortical α- synuclein pathology with a dynamic regional burden at 1-, 3-, and 6-months post-injection.
- Silver-positive neuronal processes are an early and enduring degenerative feature of the fibril model, while extensive neurodegeneration of the hippocampal CA2/3 subfield is detected at 6-months post-injection.
- Mice exhibit progressive hippocampal-dependent spatial learning and memory deficits.
- Forebrain injection of α-synuclein fibrils may be used to model aspects of Lewy-related cognitive dysfunction.

## Introduction

Lewy pathology remains an emblematic feature of neurodegenerative disease, primarily observed in Parkinson’s disease (PD) and Dementia with Lewy bodies (DLB) [1]. Having been linked to perturbations in movement and cognition, the presence of intraneuronal proteinaceous inclusions (of which α-synuclein is a key constituent) remains enigmatic [2,3]. Lewy pathology is thought to serve as a pathologic substrate for parkinsonism consequential to midbrain dopaminergic neurodegeneration. It is worth noting, however, that Lewy pathology spans well beyond the midbrain. Histopathologic schemes, such as Braak staging, have proposed that the formation of Lewy pathology follows stereotypic patterns arising first in the brainstem and advancing across the brain in a caudo-rostral manner [4,5]. Towards this, an array of experimental studies has supported a mechanism for pathologic α-synuclein spread driven by connectivity and influenced by neuronal phenotype [6]. While exceptions to this patternicity are common, forebrain structures such as the hippocampal formation are frequently affected during disease progression. It has been postulated that the spread of Lewy pathology into the forebrain may impact cognition and incite the onset of dementia [7–11]. A minority of patients with PD present with mild cognitive symptoms at baseline, and a majority (∼75%) will develop more severe deficits prior to death [12–14]. Accordingly, the relationship between Lewy pathology and cognitive dysfunction must be elucidated. Whether inclusions are indeed a driving pathologic substrate of dementia is unclear and, critically, the pathophysiology of Lewy-related cognitive dysfunction remains unresolved.

An established mouse model of “Lewy-like” α-synuclein pathology relies on the intracerebral injection of α-synuclein preformed fibrils [15–17]. Following neuronal uptake, α-synuclein fibrils recruit endogenous α-synuclein which is incorporated into neuritic and cytoplasmic inclusions, directly templating protein aggregation. These inclusions have been shown to recapitulate biochemical and histological aspects of human Lewy bodies. Importantly, the mouse α-synuclein fibril model allows for the strategic induction of pathology via stereotactic injection. Thus, the model is permissive of targeting specific circuits of interest without the developmental confounds or transcriptional artifacts present in transgenic models [18]. While the basis for neuronal vulnerability, resilience, and resistance to α-synuclein fibrils remains an area of active interest, α-synuclein fibrils have proven an effective tool to model Lewy pathology *in vivo*.

Here, we observe that α-synuclein fibril-injected mice develop impaired hippocampal-dependent spatial learning and memory as measured by the Barnes maze. Following longitudinal behavioral testing, brains were collected for histopathological assessment. Additional cohorts were used to assess the regional burden of α-synuclein pathology at 1- and 3-months post-injection (MPI) in addition to the 6 MPI behavioral cohort. As expected, α-synuclein pathology was readily induced in the mouse hippocampus and cortex with a morphology consistent with previous studies [17,19,20]. The regional burden of α-synuclein pathology was quantified at each time-point to appreciate pathological pseudo-progression. We found that the perforant pathway, composed of hippocampal-projecting neurons located in the entorhinal cortex, was especially vulnerable to the pathologic spread of α-synuclein. In contrast, the septo-hippocampal pathway, projecting from the septal region, was wholly resistant to the formation of inclusions at all time-points assessed. Concurrent with the formation of α- synuclein pathology, we observed silver-positive degenerating neuronal processes spanning both intrinsic and extrinsic hippocampal projections. By 6 MPI, we detected significant neurodegeneration in the hippocampal CA2/3 subfield. This finding was reflective of the early susceptibility of the CA2/3 subfield to α-synuclein pathology. Collectively, these pathological changes preceded the development of cognitive deficits, supporting a link between the formation of α-synuclein pathology in the forebrain and ensuing cognitive dysfunction.

## Results

### The hippocampal CA2/3 subfield exhibits early vulnerability to the formation of α- synuclein pathology

We sought to evaluate the formation of α-synuclein pathology in regions of the brain relevant to Lewy-related cognitive dysfunction. While the pathophysiological basis for cognitive dysfunction in PD and DLB is unclear, it is generally agreed that the hippocampus plays a pivotal role [21–23]. We therefore opted to target forebrain circuits relevant to spatial learning and memory in mice. Mice received bilateral stereotactic injections of either mouse α-synuclein preformed fibrils or vehicle to the hippocampus and overlaying cortex (**Fig. 1**). To assess the pseudo-progression of α-synuclein pathology over time, brains were collected and processed at 1, 3, and 6 MPI (**Fig. 1**). Coronal brain sections were subjected to immunohistochemical staining using an antibody that recognizes phosphorylated α-synuclein protein at serine 129 (pS129 α-synuclein). This is a well-established marker of misfolded α-synuclein and provides a clear indicator of inclusion pathology in our model [24]. Importantly, no discrete pS129 α-synuclein staining structures were detected in the brains of control animals supporting its use as a specific marker of pathology.

**Figure 1:**
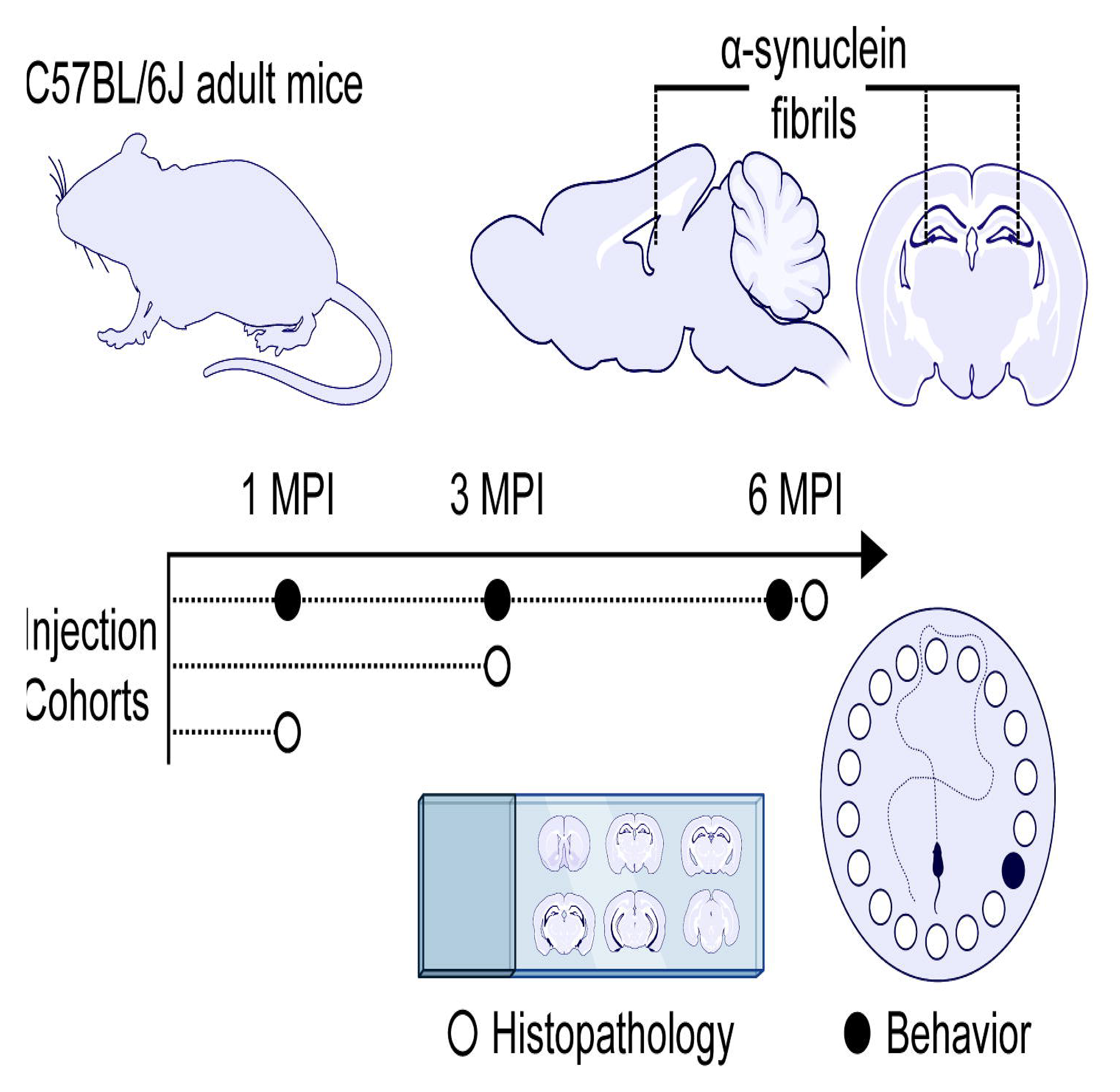
Timeline and experimental design. Adult mice (C57BL/6J) were subjected to bilateral stereotactic injection into the forebrain with either α-synuclein fibrils or vehicle. Cohort were assessed at either 1 MPI, 3 MPI, or 6 MPI.

To perform regional quantification of α-synuclein pathology, we utilized a digital slide scanning system (Aperio) paired with HALO analysis software (Indica Labs). This facilitated on-slide digital annotation and uniform quantitative assessment of α-synuclein pathology at the regional level across tissue-sections and time-points. We opted to focus our analysis on selected forebrain structures based on the appreciable pattern of pS129 α-synuclein immunostaining. The robust induction of α-synuclein pathology was observed in the hippocampus, including the dentate gyrus and CA1-CA3 subfields, with clear distinctions in regional burden across the assessed time-points (**Fig. 2A-D**). Cortical pathology was detected to a lesser extent (**Fig. S1A-C**). We therefore elected to focus much of our analyses on spatiotemporal differences in pathology burden in the hippocampal subfields.

**Figure 2:**
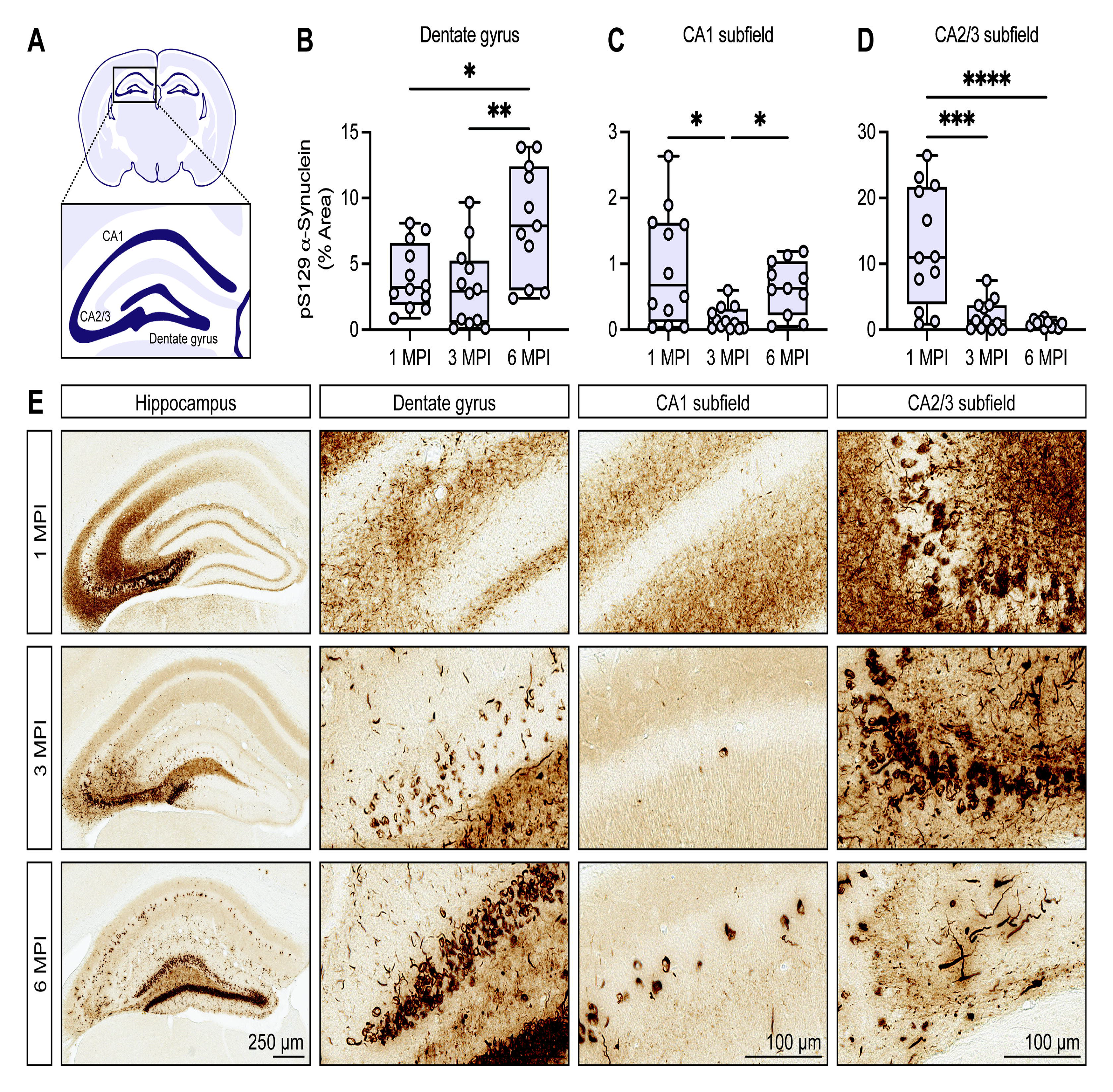
Formation of α-synuclein pathology in the hippocampal subfields. **A)** Schematic representation of the mouse hippocampal subfields, including the dentate gyrus, CA1 and CA2/3 subfields. **B-D)** Digital pathology quantification of pS129 α- synuclein immunostaining in the **(B)** dentate gyrus, **(C)** CA1 subfield, and **(D)** CA2/3 subfield at 1, 3, and 6 MPI. Data are expressed as boxplots depicting the median, interquartile range, and individual data points of the % area occupied of pS129 α- synuclein immunostaining (*n* = 11-12 animals/group/timepoint). Dentate gyrus, CA2/3 subfield: \**P*<0.05, \*\**P*<0.01, \*\*\**P*<0.001, or \*\*\*\**P*<0.0001 by one-way ANOVA with Tukey’s multiple comparisons test, as indicated. CA1 subfield: \**P*<0.05 by Kruskal-Wallis test with Dunn’s multiple comparisons test, as indicated. **E)** Representative images of pS129 α-synuclein immunostaining at 1, 3, and 6 MPI including the dorsal hippocampus and each subfield. Scale bar: 250 μm (hippocampus) or 100 μm (dentate gyrus, CA1 subfield, CA2/3 subfield).

To better understand the pseudo-progression of α-synuclein pathology in our model, we assessed hippocampal subfield burden at 1, 3, and 6 MPI (**Fig. 2A-D**). At 1 MPI, tendril-like somatic inclusions were observed in the hippocampal CA2/3 subfield while immunostaining in the other subfields was predominantly neuritic (**Fig. 2E**). Extensive neuritic pathology was also observed adjacent to the CA2/3 pyramidal layer at 1 MPI, which appeared to coalesce towards somatic inclusions at later time-points (**Fig. 2E**). This is consistent with previous studies exhibiting relatively early peak inclusion formation with a reduction in burden over time [25–27]. To be clear, neuritic pathology was still observed across all time-points but was greatly reduced by 3 MPI. In parallel, an increased burden of somatic inclusions was observed in the dentate gyrus and CA1 subfield over time (**Fig. 2B-C**). In contrast, the burden of α-synuclein pathology in the CA2/3 hippocampal subfield was drastically reduced at later time-points (**Fig. 2D**). This was at least partially related to the decline in diffuse immunostaining after 1 MPI but was also characterized by an obvious decrease in cytoplasmic inclusions in portions of the pyramidal layer at 6 MPI (**Fig. 2E**).

### Hippocampal projections arising from the entorhinal cortex are vulnerable to pathologic **α**-synuclein spread while projections from the medial septal area are resistant

Distal from the injection site, we observed a considerable burden of α-synuclein pathology in the entorhinal cortex. This is consistent with retrograde spread of pathology via entorhinal-hippocampal projections, which comprise the perforant pathway (**Fig. 3A**). We observed the formation of dense cytoplasmic inclusions along with neuritic pathology at 1 MPI, with a reduction in the burden of pathology at subsequent time-points (**Fig. 3B-C**). While we did not assess α-synuclein pathology to the level of each cortical layer, pyramidal neurons in layers II/III were predominantly affected. This may be partially explained by established patterns of entorhinal-hippocampal connectivity, with entorhinal layers II/III projecting to distinct targets from entorhinal layers IV/V [28]. Alternatively, layers II/III may be especially vulnerable to inclusion formation due to cell autonomous factors, such as higher endogenous levels of α-synuclein. Previous studies support that endogenous α-synuclein protein levels are a strong determinant of inclusion formation in the α-synuclein fibril model [26,29].

**Figure 3:**
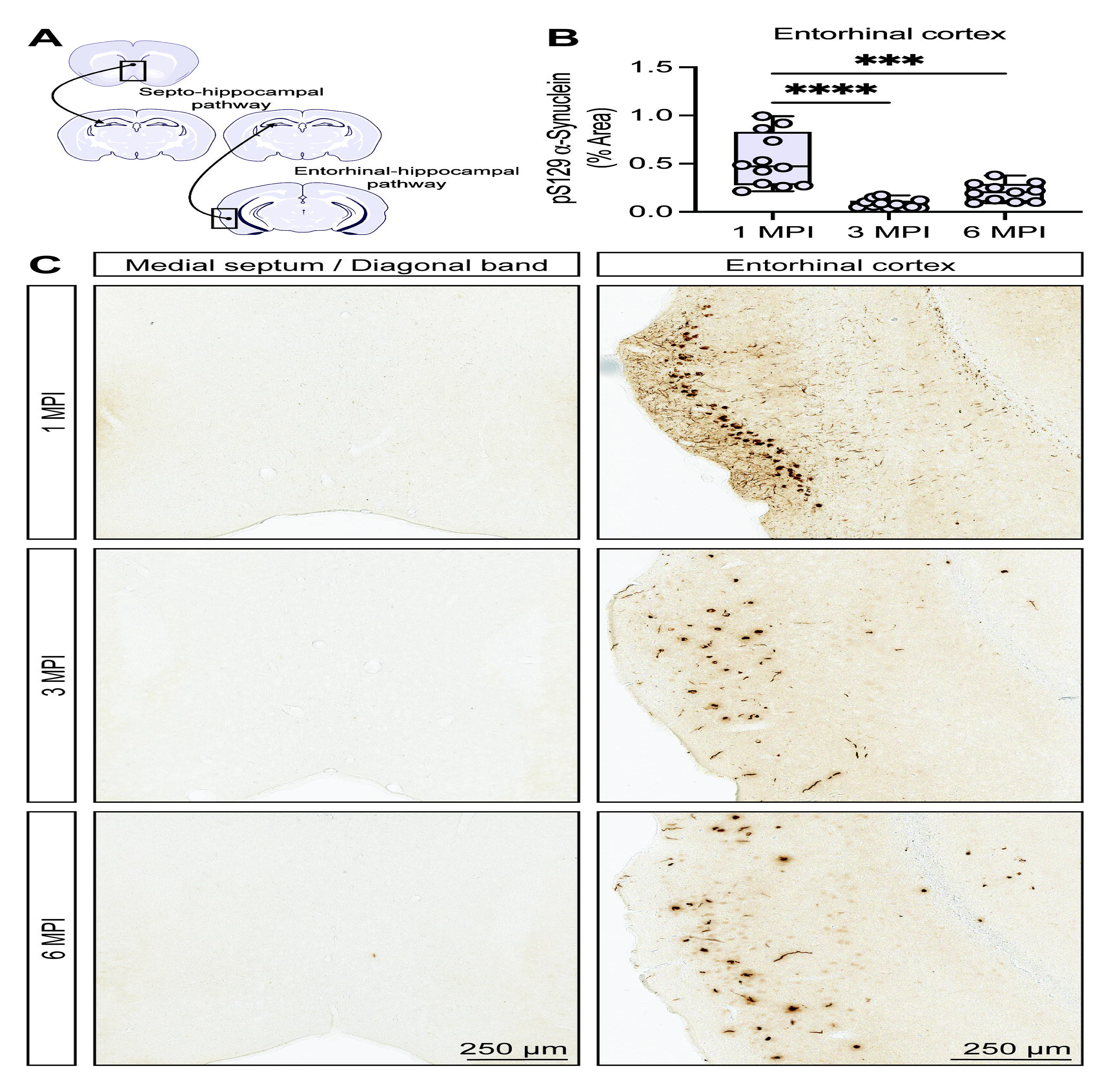
Divergent susceptibility of hippocampal projections to pathologic spread of α-synuclein pathology. **A)** Schematic representation of the septo-hippocampal and entorhinal-hippocampal pathways with neuronal populations labeled by a dot and a line with arrow projecting to the hippocampus. **B)** Digital pathology quantification of pS129 α-synuclein immunostaining in the entorhinal cortex at 1, 3, and 6 MPI. Data are expressed as boxplots depicting the median, interquartile range, and individual data points of % area occupied of pS129 α-synuclein immunostaining (*n* = 11- 12 animals/group/timepoint). \*\*\**P*<0.001 or \*\*\*\**P*<0.0001 by one-way ANOVA with Tukey’s multiple comparisons test, as indicated. **C)** Representative images of pS129 α- synuclein immunostaining at 1, 3, and 6 MPI in the medial septum/diagonal band (left column) and entorhinal cortex (right column). Scale bar: 250 μm.

In contrast to the entorhinal-hippocampal pathway, medial septal neurons in the septo-hippocampal pathway did not exhibit cytoplasmic inclusions at any time-point assessed (**Fig. 3C**) aside from the occasional labeled neurite. The septo-hippocampal pathway is known to degenerate in some neurodegenerative mouse models, and previous studies using injection of tau paired helical filaments in the hippocampus have resulted in transmission of tau pathology to the medial septal area and diagonal band [30–32]. Intriguingly, this does not appear to be the case with α-synuclein fibrils. Consistent with our findings, a recent study examining α-synuclein fibril injection also failed to detect pathology in this region, despite the detection of a co-injected tracer [33]. It is possible that a greater duration of time is required for the spread of pathology into this region. In support of this premise, a study characterizing fibril injection in rats found that the vertical limb of the diagonal band developed pathology at later time-points relative to hippocampal and cortical areas [34]. Much interest has been placed on the trans-neuronal spread of α-synuclein fibrils, while less emphasis has been spent on identifying cell autonomous mechanisms of susceptibility. Given our limited understanding of neuronal vulnerability, the basis for this resilience in the basal forebrain may be particularly informative and worth further exploration in fibril models.

### Silver-positive degenerating neuronal processes are observed in regions affected by **α**-synuclein pathology

Previous studies have demonstrated that exposure to α-synuclein fibrils results in alterations to spine morphology accompanied by perturbed synaptic activity [29,35,36]. This supports that axonal damage may be an early pathologic feature caused by the corruption of endogenous α-synuclein following exposure to α-synuclein fibrils. Moreover, axonal degeneration is a key feature of PD and is speculated to contribute to disease pathogenesis [37]. To examine whether similar degenerative changes were present in our model, we performed Gallyas silver staining and examined pathologically affected regions. Since we observed neuritic α-synuclein pathology in the supra-pyramidal region of the CA1 subfield at 1 MPI, we sought to examine whether Schaffer collaterals might exhibit signs of degeneration (**Fig. 4A-B**). Likewise, given the spread of α-synuclein pathology from the hippocampus to the entorhinal cortex, we also considered the integrity of projections in the perforant pathway (**Fig. 4A-B**). Both regions revealed marked silver-positive degenerating neuronal processes at all assessed time-points, suggestive of early and persistent axonal damage following fibril injection (**Fig. 4C**). As expected, no evidence of silver-positive neurite degeneration was observed in control animals (**Fig. 4C**).

**Figure 4:**
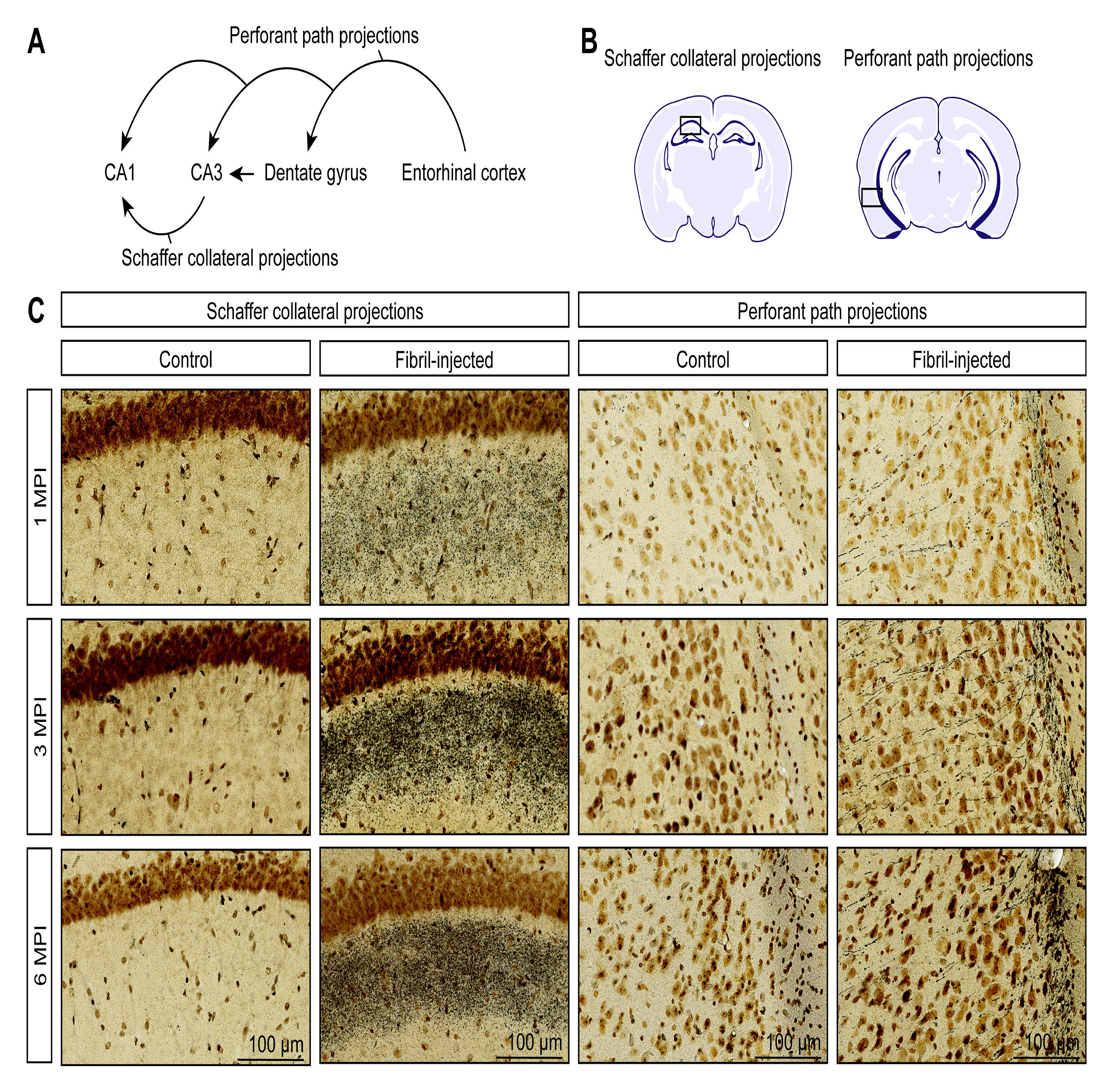
Evidence of axonal damage concurrent with the formation of α-synuclein pathology. **A)** Schematic representation of the “tri-synaptic circuit”, depicting the perforant path and Schaffer collaterals spanning the entorhinal cortex and hippocampus. **B)** Schematic representation of the regions of interest depicted in histology images **(C)**: the CA1 subfield and the entorhinal cortex. **C)** Representative images of Gallyas Silver staining (gray/black fibers or puncta) at 1, 3, and 6 MPI in fibril-injected mice. Data are representative *n* = 4 animals/group/timepoint. Scale bar: 100 μm.

### Extensive neuronal loss is observed in the pyramidal layer of the hippocampal CA2/3 subfield following the formation of **α**-synuclein pathology

Since a clear reduction in the burden of α-synuclein pathology was observed within the hippocampal CA2/3 subfield, we sought to determine whether this reduction might be explained by neuronal loss. It is likely that an initial drop in α-synuclein pathology was at least partially related to sequestration of pathologic inclusions to the neuronal perikaryon. This plausibly explains the drastic reduction from 1 MPI to 3 MPI, as neuritic pathology held a considerable fraction of immunostaining area at 1 MPI (**Fig. 2D-E**). By 6 MPI, however, there was an appreciable reduction in cytoplasmic inclusions in the portion of the pyramidal layer that exhibited inclusions at earlier time-points. We next evaluated a broadly expressed neuronal nuclei protein (NeuN) to probe for neuronal loss within the hippocampal subfields. We found that fibril-injected mice exhibit a marked reduction (∼50%) in NeuN-positive immunostaining within the CA2/3 subfield at 6 MPI, consistent with neuronal loss (**Fig. 5C-D**). While we did not assess hippocampal atrophy, this reduction was accompanied by observable atrophy of the CA2/3 subfield which further supported that loss of NeuN-positivity was due to neurodegenerative changes (**Fig. 5D**). In contrast, we did not find evidence of significant neuronal loss in either the dentate gyrus or CA1 subfield of fibril-injected mice despite the presence of α- synuclein pathology (**Fig. 5A-B**). Given that glia are known to contribute to neurodegenerative processes, we also surveyed for astroglial reactivity as measured by GFAP immunostaining [38]. While not a central focus of this study, we detected increases in astroglial reactivity over time, particularly in the CA2/3 subfield at 3 MPI and across multiple regions at 6 MPI (**Fig. S5A-D**).

**Figure 5:**
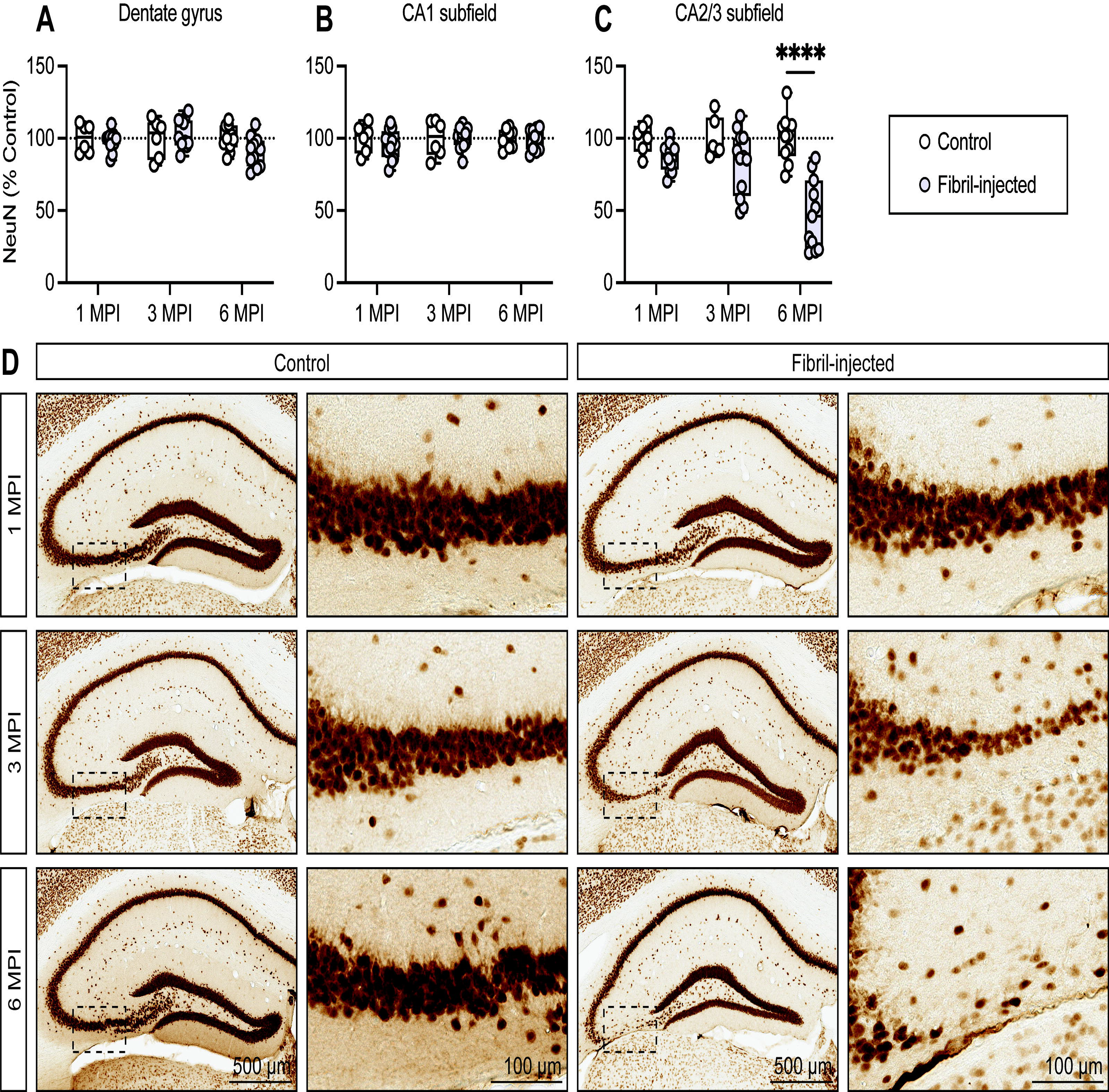
Neuronal loss is observed in the CA2/3 hippocampal subfield. **A-C)** Quantification of NeuN-positive immunostaining in the **(A)** dentate gyrus, **(B)** CA1 subfield, and **(C)** CA2/3 subfield. Data are expressed as boxplots depicting the median, interquartile range, and individual data points for NeuN-positive area normalized to control at 1, 3, and 6 MPI in the **(A)** dentate gyrus, **(B)** CA1 subfield, and **(C)** CA2/3 subfield (*n* = 6-12 animals/group/timepoint). \*\*\*\**P*<0.0001 by two-way ANOVA with Bonferroni’s multiple comparisons test, as indicated. **D)** Representative images of NeuN immunostaining in the hippocampus and CA2/3 subfield in control (left columns) and fibril-injected mice (right columns) at 1, 3, and 6 MPI. Scale bar: 500 μm (hippocampus) or 100 μm (CA2/3 subfield).

### Mice exhibit increasingly impaired performance in a hippocampal-dependent cognitive task following injection with **α**-synuclein fibrils

Studies exploring the use of α-synuclein fibrils to model PD *in vivo* have generally adopted an intrastriatal paradigm [26,27,39–44]. Several of these studies have demonstrated the onset of motor deficits following the formation of α-synuclein pathology in the substantia nigra along with dopaminergic neurodegeneration. We hypothesized that a similar paradigm applying α-synuclein fibrils to the forebrain might be sufficient to elicit cognitive dysfunction.

To examine general motoric behavior at 6 MPI, we employed open-field testing which fails to reveal significant differences between experimental groups (**Fig. S6C**). Similarly, we did not detect any influence of α-synuclein fibril-injection on anxiety-related phenotypes in the elevated plus maze or marble-burying test at 6 MPI (**Fig. S6A, D**). Executive function appears to be unaffected as well, as no observable changes were seen in the nest-building assay (**Fig. S6B**). In general, we did not perceive any indications of motor dysfunction and, as expected, the midbrain was devoid of α- synuclein pathology in fibril-injected mice (data not shown).

Following injection of either α-synuclein fibrils or vehicle, mice were longitudinally assessed using the Barnes radial maze at 1, 3, and 6 MPI. We utilized this assay to probe spatial learning and memory performance over time (**Fig. 6A**). Importantly, performance in the Barnes maze is hippocampal-dependent and improvement during acquisition trials, as measured by latency, is indicative of spatial learning [45–47]. At 1 MPI, we did not detect a significant difference in performance between experimental groups suggesting the absence of cognitive dysfunction in fibril-injected mice at our earliest time-point (**Fig. 6B** and **Fig. S7A**). This is intriguing, as α-synuclein pathology is seemingly abundant at 1 MPI (**Fig. 2E**). Assessment of fibril-injected mice at later time-points revealed a progressive performance deficit relative to control mice, as measured by average performance as well as across acquisition trials (**Fig. 6C-D** and **Fig. S7B-C**). Intriguingly, spatial reference memory was unaffected by α-synuclein fibril injection at either 3 or 6 MPI (**Fig. 6H-I**). This suggests that fibril-injected mice retain some capacity for spatial memory, but that aspects of learning may be limited.

**Figure 6:**
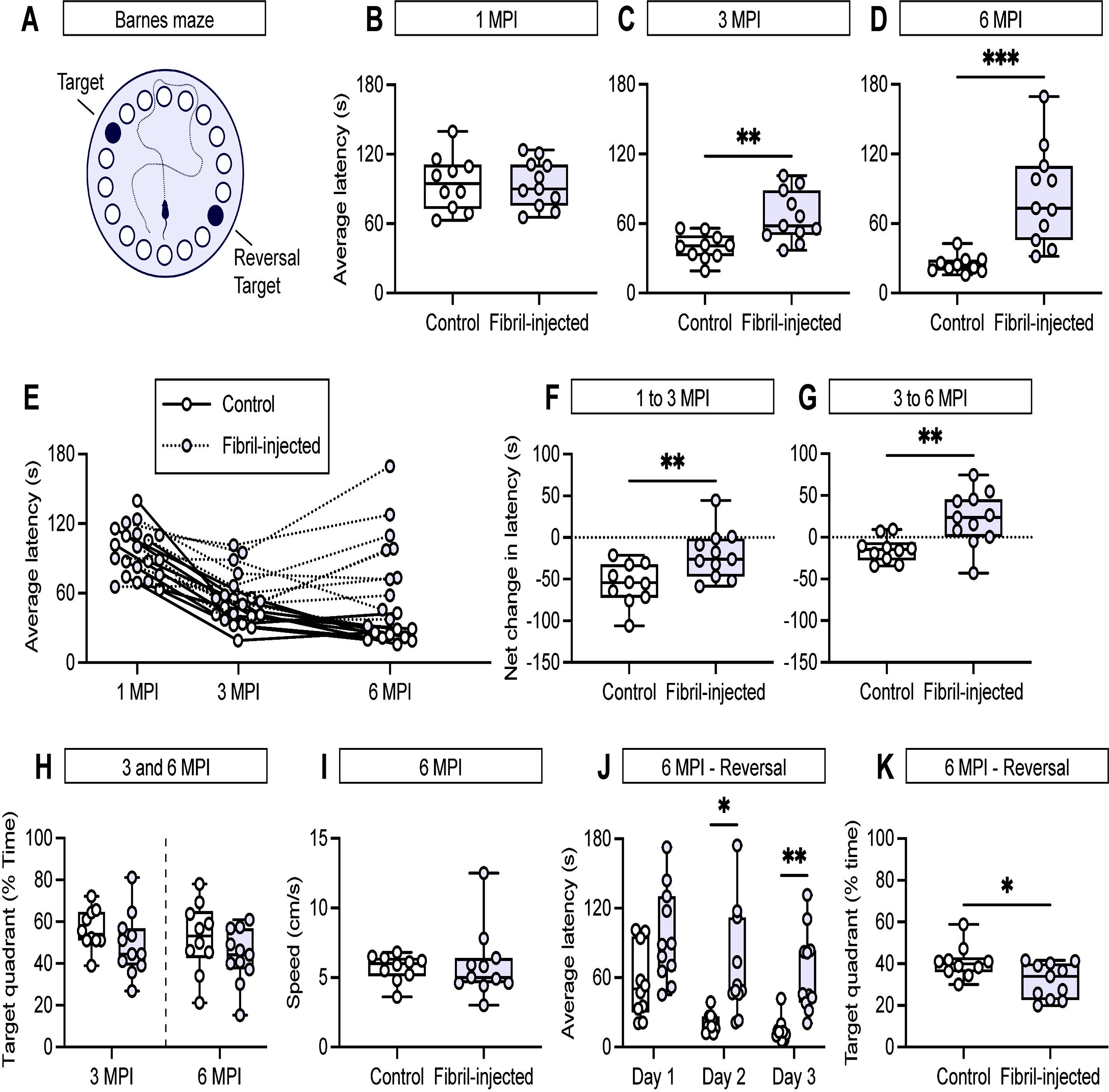
Mice injected with α-synuclein fibrils develop cognitive dysfunction. **A)** Schematic representation of the Barnes maze platform in which the target and reversal target are 180° apart. **B-D)** Quantification of the average latency across acquisition trials at **(B)** 1 MPI, **(C)** 3 MPI, and **(D)** 6 MPI. Data are expressed as boxplots depicting the median, interquartile range, and individual data points for the average latency of control and fibril-injected mice (*n* = 10-11 animals/group). \*\**P*<0.01 or \*\*\**P*<0.001 by unpaired Student’s *t*-test. **E)** Spaghetti plot depicting the average latency of mice at 1, 3, and 6 MPI. Each line represents the performance trajectory of an individual mouse. **F-G)** Quantification of performance trajectory from **(F)** 1 MPI to 3 MPI and **(G)** 3 MPI to 6 MPI. Data are expressed as boxplots depicting the individual data points for the net change in average latency of each group from 1 MPI to 3 MPI and from 3 MPI to 6 MPI. \*\**P*<0.01 by unpaired Student’s *t*-test. **H)** Boxplots depicting individual data points for the percentage of time in which mice occupied the target quadrant during the probe session. **I)** Boxplots depicting the individual data points for the average speed during the probe session. **J)** Boxplots depicting individual data points for the average latency of each group during the reversal segment of the Barnes maze. \*\**P*<0.01 or \*\*\**P*<0.001 by two-way ANOVA with Bonferroni’s multiple comparisons test, as indicated. **K)** Boxplots depicting individual data points for the percentage of time in which mice occupied the target quadrant during the probe session during the reversal segment of the Barnes maze. \**P*<0.05 by unpaired Student’s *t*-test.

At 6 MPI, we also utilized a reversal paradigm (relocating the placement of the target hole by 180°) (**Fig. 6A**). Under the reversal paradigm, fibril-injected mice again exhibited impaired spatial learning relative to control mice (**Fig. 6J**). In addition, fibril-injected mice spent less time in the target quadrant during the reversal probe session, indicating that the additional cognitive load of the reversal paradigm was sufficient to impair spatial memory (**Fig. 6K**).

To better characterize the impact of fibril-injection on longitudinal trends in the Barnes maze, we evaluated the performance trajectory of each mouse. As expected, mice in the control group demonstrated reliable improvement in the Barnes maze across successive time-points, supportive of increased familiarity with the assay and intact spatial cognition (**Fig. 6E-G**). Notably, fibril-injected mice failed to improve under the same conditions and generally exhibited a worse average performance at 6 MPI (**Fig. 6E-G**). These findings point towards a compromised cognitive trajectory in fibril-injected mice.

Given our interest in whether the formation of α-synuclein inclusions would be sufficient to cause cognitive dysfunction, we further analyzed the relationship between α-synuclein pathology, hippocampal neurodegeneration, and Barnes maze performance at 6 MPI. Interestingly, we find that NeuN-positive neuronal loss in the CA2/3 subfield was associated with increased levels of residual α-synuclein pathology at both 3 and 6 MPI (**Fig. S2F, I**). We identified a similar finding in the dentate gyrus at 6 MPI (**Fig. S2G**), but no other significant associations were present (**Fig. S2A-E, H**). In terms of behavior, poorer Barnes maze performance during the 6 MPI acquisition phase was significantly associated with NeuN-positive neuronal loss in the CA2/3 subfield, but not with any regional α-synuclein burden (**Fig. S3E**). In addition, we assessed the relationship between neurodegeneration in the CA2/3 subfield and the formation of pathology in the adjacent subfields and entorhinal cortex. We detected a significant association between neuronal loss in the CA2/3 subfield with greater α-synuclein burden in the dentate gyrus and CA1 subfield, but not the entorhinal cortex (**Fig. S4A-C**). Thus, is appears that neurodegeneration may directly or indirectly potentiate the spread of pathology. While the mechanism(s) underlying the pathophysiology of Lewy-related cognitive dysfunction remain to be further elucidated, our results support α- synuclein inclusions as a primary neuropathological substrate. However, neurodegeneration rather than α-synuclein burden may be more predictive of cognitive deficits.

## Discussion

Here, we sought to evaluate the formation of α-synuclein pathology across forebrain structures following fibril-injection, and whether this might precipitate cognitive impairment. A striking observation of this study was the temporally dynamic pattern of α- synuclein pathology within the hippocampus and cortex. While some inter-mouse variability was present, the pattern of distribution within the hippocampal formation remained markedly consistent with the CA2/3 subfield exhibiting the greatest vulnerability to pathology. This finding is comparable with recent estimates of Lewy pathology in the human hippocampus, which identify the CA2/3 subfield as a heavily affected subfield in Lewy body disease subjects [9,11,48–50]. We also identified a septal-hippocampal input with relative resistance to pathology, in contrast to what has been observed in human disease [7,51]. Our model is therefore able to recapitulate at least some, but not all, aspects of the regional distribution of Lewy pathology in the forebrain. Targeting hippocampal circuits via alternative injection paradigms might further clarify the relationship between connectivity and regional vulnerability, aside from what might be explained by endogenous levels of α-synuclein.

In addition to characterizing pathologic spread in our model, we also identified accompanying neurodegenerative features. These findings are consistent with previous studies demonstrating that select hippocampal neuronal populations exhibit greater vulnerability to α-synuclein inclusion formation and toxicity [19]. Towards this, we identified silver-positive degenerative processes both within the hippocampus proper and arising from the entorhinal cortex. This supports that the uptake of fibrils and resultant corruption of endogenous α-synuclein is ultimately harmful to neurons within these affected regions. In support of this notion, we observed substantial loss of NeuN-positive pyramidal neurons in the hippocampal CA2/3 subfield at 6 MPI, the same population that exhibited early formation of cytoplasmic inclusions. Thus, degenerative neuronal processes appear to be an early feature of our model and may indicate a proclivity towards neurodegeneration. While we did not separate pS129 α-synuclein immunostaining localizing to neuronal perikarya or neurites, the affected cellular compartment may be an important aspect of disease pathobiology, with relevance to therapeutic development. Accordingly, future studies in this model would be enhanced by a neuronal compartment-specific dissection of α-synuclein pathology.

Notably, we did not detect a cognitive deficit during our earliest behavioral assessment. Fibril-injected mice performed comparably to controls at 1 MPI despite indications of a substantial α-synuclein burden and axonal damage during this period. At later timepoints, and especially in the presence of neuronal loss, we detected a compromised performance in the Barnes maze. This deficit was specific to spatial learning and memory, as other assessments of motor and anxiety-related changes were unremarkable. Importantly, these results build upon previous efforts to define the influence of α-synuclein inclusion pathology on non-motor phenotypes [52–54]. Under a similar paradigm, a recent study reported hippocampal fibril-injection was insufficient to produce cognitive deficits at 3 MPI [33]. The consistent absence of cognitive dysfunction in the presence of α-synuclein pathology suggests that inclusions alone are insufficient to produce functional deficits. Towards this, neuronal loss was not detected in their model [33]. In contrast, a separate study found that α-synuclein fibrils were able to induce α-synuclein pathology and working memory deficits as early as 8-weeks, with the notable detection of cell death in the hippocampus [35]. Aside from fibril-based models, transgenic α-synuclein models have been shown to develop age-dependent cognitive phenotypes, though the co-occurrence with neurodegeneration is mixed and it is unclear the extent to which α-synuclein overexpression impacts the development of functionally relevant circuits [56].

To delineate the impact of α-synuclein pathology more clearly, we applied α- synuclein fibrils to the forebrain and utilized a hippocampal-dependent behavioral paradigm. Our results suggest that the mere presence of α-synuclein pathology is insufficient to produce cognitive deficits shortly following inclusion formation. An obvious caveat of our study is that we did not conduct a more detailed characterization of α- synuclein inclusions over time. While pS129 α-synuclein is a validated histological marker of aggregated α-synuclein, we did not assess biochemical changes that would be undetected by our approach. Given that neurons can harbor pathological inclusions over substantial timeframes, the evolution of α-synuclein pathology within an individual neuron may be an important factor to consider. It may be the case that a “pathologic threshold” must be reached prior to neuronal dysfunction.

Our study examined α-synuclein pathology in the context of hippocampal circuits, but it is important to consider that other circuits have been implicated in the non-motor behavioral spectra of PD and DLB as well. A recent study found that intrastriatal injection of α-synuclein fibrils in mice led to extensive inclusion formation in the cortex and amygdala [52]. Under this paradigm, deficits in social dominance and fear conditioning were detected in the absence of neurodegeneration at 6 MPI. These results suggest that neuronal loss is not an absolute requisite for the manifestation of cognitive dysfunction, though it is unclear whether neurodegeneration might have been detected in their model given a longer timeframe. An incubation period, of sorts, may therefore be required before the formation of inclusions precipitates detectable behavioral changes. In our model, we were unable to clearly dissociate whether α- synuclein pathology ultimately contributes to cognitive phenotypes independent of neuronal loss, although we did detect behavioral changes in the absence of significant neurodegeneration at 3 MPI. This question should be further explored, as it is worth determining whether α-synuclein contributes to cognitive changes in a manner converging with, or in parallel to, overt neurodegeneration. Importantly, we observed silver-positive degenerating neuronal processes early in our model. This finding suggest that pathological inclusions have an immediate toxic effect in at least some neuronal populations. Further, degenerating neuronal processes may be highly relevant to cognitive changes, as α-synuclein pathology may disrupt circuit function prior to cell death.

The hippocampal formation has been implicated in Lewy-related cognitive dysfunction, with antemortem memory performance correlating with α-synuclein pathology across hippocampal subfields and the entorhinal cortex [9]. While additional studies have supported a link between forebrain Lewy pathology and cognitive status, the degree to which α-synuclein drives cognitive dysfunction is unclear. Nonetheless, nearly all patients will experience some level of cognitive impairment [12]. Accordingly, we tested the hypothesis that forebrain injection of α-synuclein fibrils would be sufficient to induce cognitive dysfunction in mice. Our findings support that the formation of forebrain α-synuclein pathology leads to behavioral deficits consistent with the complex cognitive features observed in PD and DLB.

## Materials and methods

### Animals

We used 10- to 12-week-old wild type C57BL/6J mice obtained from the VAI Vivarium internal colony. Both male and female mice were used for this study. Mice were housed at a maximum of 4 per cage under a 12-hour light/dark cycle and were provided food and water *ad libitum*. Animal studies were reviewed and approved by the Institutional Animal Care and Use Committee at Van Andel Institute. Mice were maintained in accordance with NIH guidelines for care and use of animals.

### Preparation of mouse **α**-synuclein fibrils

Preparation of α-synuclein fibrils from wild-type, full-length, recombinant mouse α- synuclein monomers was completed as previously reported [20,57,58]. Prior to surgeries, fibrils were thawed to room temperature and sonicated for a period of 4 min in a Q700 cup horn sonicator (Qsonica) at 50% power for 120 pulses (1 s on, 1 s off). Fibrils used for injection were diluted to a concentration of 5 μg/μl.

### Stereotactic surgeries

Mice were anesthetized with 2% isoflurane anesthesia and positioned within a stereotactic frame (Kopf). Mice received bilateral injections using the following coordinates relative to bregma: anterior-posterior (A-P), −2.5 mm; medio-lateral (M-L), +/-2.0 mm; dorso-ventral (D-V), −2.4, −1.4 mm. Each injection was delivered in a volume of 1 μl at a flow rate of 0.2 μl/min. Animals were sacrificed at 1-, 3-, or 6-months post-injection.

### Behavioral analyses

Behavioral studies occurred in a mouse behavioral testing suite using ANY-maze software (Stoelting) in conjunction with a ceiling-mounted video camera to track and record behavior during testing. Fibril-injected and control mice were randomly intermixed during testing in a blinded manner. Prior to any testing, mice were habituated to handling and allowed to acclimate to the testing suite. All behavioral studies that did not require testing on consecutive days were interspaced to provide animals with a minimum 24 h period of rest. Assays which required repeated testing, or in which mice were placed in the same apparatus sequentially, were subjected to rigorous inter-trial cleaning with 70% ethanol. Standard parameters for all assays were maintained and followed across longitudinal testing periods.

### Open-field test

Mice were placed in the open-field apparatus for a 20 min assessment period. Several metrics regarding mouse behavior were quantified including average speed, total distance travelled, and time spent in the center versus the periphery of the apparatus.

### Marble-burying test

Mice were individually placed in a standard housing bin filled with 4x the typical amount of bedding material, along with 20 marbles (1 cm diameter, clear) arranged in a 4 x 5 array on the surface of the bedding material. Following a 30 min period, mice were gently removed from the housing bin and each bin was photographed for later scoring. Images were scored for the number of visible marbles across the surface of the bedding material. A calculation was then performed to determine the number of buried marbles.

### Elevated-plus maze

Mice were placed in the enclosed corner of the elevated-plus maze apparatus at the start of the assay. Mice were allowed to freely explore the apparatus over a 10 min period. Time spent in either the enclosed or open arms was determined along with the number of total arm entries. The apparatus was thoroughly cleaned with 70% ethanol between animals.

### Barnes maze

The Barnes maze was performed as previously described with some modifications [46]. A white Barnes maze platform with 20 holes spaced equi-distant along the outer perimeter was used for all paradigms during the study. An acquisition phase of 4 trials per day was followed for a total of 4 days (12-16 total trials per animal). The maximum trial duration was limited to 180 s with a 20 min inter-trial interval. The duration for the animals’ head to enter the target hole (latency) during each trial was evaluated using ANY-maze software. The assay was repeated at 3 MPI and again at 6 MPI, with the addition of a probe phase occurring 24 h following the final acquisition phase trial sessions. At 6 MPI, the probe phase was also followed by reversal acquisition and probe phases in which the location of the target hole was rotated 180°.

### Immunohistochemistry

Mice were transcardially perfused with 4% paraformaldehyde (PFA) in 0.1 M phosphate buffer (pH 7.4). Brains were removed, post-fixed for 24 hours in 4% PFA and cryoprotected in 30% sucrose solution, and 35 μm-thick coronal sections were prepared. For immunostaining, sections were quenched for endogenous peroxidase activity by incubation in 3% H_2_O_2_ (Sigma) diluted in methanol for 5 min at 4°C. Sections were then blocked in 10% normal goat serum (Invitrogen), 0.1% Triton-X100 in PBS for 1 hour at room temperature. Sections were then incubated with primary antibodies for 48 hours at 4°C followed by incubation with biotinylated secondary antibodies (Vector Labs) for 24 hours at 4°C. Following incubation with ABC reagent (Vector Labs) for 1 hour at room temperature and visualization in 3, 3’-diaminobenzidine tetrahydrochloride (DAB; Vector Labs), sections were mounted on Superfrost plus slides (Fisher Scientific). Slide-mounted sections were then dehydrated with increasing ethanol concentrations and xylene, and coverslipped using Entellan (Merck). The following primary antibodies were used: rabbit anti-pS129-α-synuclein (ab51253; Abcam), mouse anti-NeuN (MAB377; Sigma-Aldrich), rabbit anti-GFAP (ab227761; Abcam). The following biotinylated secondary antibodies were used: goat anti-mouse IgG and goat anti-rabbit IgG (Vector Labs).

### Quantitative pathology analyses

Digital scans of slide-mounted coronal sections at 20x magnification were generated using a ScanScope XT slide scanner (Aperio) at a resolution of 0.5 µm/pixel. Quantitative analyses were performed using HALO analysis software (Area quantification and Object colocalization modules; Indica Labs Inc.) as previously described [59]. Threshold parameters were developed within the modules to provide broad detection of markers without the inclusion of background staining. Anatomical regions of interest were annotated across a minimum of 3 to 5 coronal sections using the Allen Mouse Brain Atlas (Allen Institute) as a guide [60]. For specific quantification of NeuN staining in the hippocampus, the pyramidal cell layer of the CA1 and CA2/3 subfields, and granule cell layer of the dentate gyrus were manually outlined to measure total immunostained area. NeuN-positive area was then normalized to generate a % NeuN-positive area relative to controls. All sections/images were batch analyzed in Halo analysis software using the developed parameters.

### Statistical analyses

Boxplots depict the minimum to maximum data points with median and interquartile range. All data was tested for normality using a Shapiro-Wilk test. Data that fit a normal distribution was analyzed using either an independent two-tailed unpaired Student’s *t*- test, one-way ANOVA, or 2-way ANOVA. Data that was not normally distributed was analyzed using a Kruskal-Wallis test. *P* values were represented as follows: * *P*<0.05, \*\**P*<0.01, \*\*\**P*<0.001, \*\*\*\**P*<0.0001. Graphs were generated with GraphPad Prism 9 (GraphPad Software).

## Declarations

### Ethical approval and consent to participate

All animal procedures were conducted in accordance with the guidelines set forth by the Institutional Animal Care and Use Committee (IACUC) at Van Andel Institute, and study protocols were reviewed and approved prior to performing the experimental procedures described.

## Consent for publication

All authors read and approved the final manuscript.

## Availability of supporting data

All data generated or analyzed during this study are included in this published article and its supplementary information.

## Competing interests

The authors declare they have no financial competing interests.

## Funding

This work was supported by a grant from the National Institutes of Health (NIH) R56AG074473 to DJM, and by financial support from the Van Andel Institute and the Van Andel Institute Graduate School.

## Author contributions

DJM conceptualized and supervised the research; DJD planned and executed most experiments, contributed to research conception, design and interpretations; DJD wrote the manuscript with the help of DJM; APN contributed to research design and interpretation; KB and JM produced and validated α-synuclein preformed fibrils. All authors read and approved the final manuscript.

## Acknowledgements

We thank the VAI Vivarium, Optical Imaging, Biostatistics, and Pathology and Biorepository Cores for technical assistance. Some figures were created using BioRender.com.

## Supplementary Figure Legends

**Figure S1: Formation of α-synuclein pathology in the cortex. A)** Schematic representation of the region of interest within the mouse cortex. **B)** Digital pathology quantification of pS129 α-synuclein immunostaining in the cortex at 1, 3, and 6 MPI. Data are expressed as boxplots depicting the median, interquartile range, and individual data points of % area occupied of pS129 α-synuclein immunostaining (*n* = 11-12 animals/group/timepoint). Non-significant by Kruskal-Wallis test with Dunn’s multiple comparison test. **C)** Representative images of pS129 α-synuclein immunostaining at 1, 3, and 6 MPI in the cortex. Scale bar: 250 μm.

**Figure S2: Relationship between neuronal loss and residual α-synuclein pathology in the hippocampus. A-I)** Association between normalized NeuN-levels and pS129 α-synuclein immunostaining in the **(A, D, G)** dentate gyrus, **(B, E, H)** CA1 subfield, and **(C, F, I)** CA2/3 subfield at **(A, B, C)** 1 MPI, **(D, E, F)** 3 MPI, and **(G, H, I)** 6 MPI (*n* = 11-12 animals/group). Data are expressed as individual data points. \**P*<0.05 by Spearman correlation, *r* values listed by region, as indicated.

**Figure S3: Cognitive performance is associated with neuronal loss in the CA2/3 subfield but not residual α-synuclein burden. A, C, E)** Association between average latency in the Barnes maze at 6 MPI and (A, C, E) NeuN-levels in the (A) dentate gyrus, (C) CA1 subfield, and (E) CA2/3 subfield in the 6 MPI cohort. \*\**P*<0.01 by Spearman correlation, *r* values listed by region, as indicated. B, D, F) Association between average latency at 6 MPI and (B, D, F) pS129 α-synuclein immunostaining in the (B) dentate gyrus, (D) CA1 subfield, and (F) CA2/3 subfield at 6 MPI. Not significant by Spearman correlation, *r* values listed by region, as indicated.

**Figure S4: Spread of α-synuclein pathology into the dentate gyrus and CA1 subfield is associated with neuronal loss in the CA2/3 subfield. A-C)** Association between CA2/3 subfield NeuN-levels and pS129 α-synuclein immunostaining in adjacent regions at 6 MPI including **(A)** dentate gyrus, **(B)** CA1 subfield, and **(C)** entorhinal cortex. \**P*<0.05 by Spearman correlation, *r* values listed by region, as indicated.

**Figure S5: Astrogliosis is detected in fibril-injected mice. A)** Representative images of GFAP immunostaining at 1, 3, and 6 MPI in the hippocampal subfields and cortical regions, arranged by fibril-injected and control group. **B-D)** Digital pathology quantification of GFAP immunostaining by region at 1, 3, and 6 MPI. Data are expressed as boxplots depicting the median, interquartile range, and individual data points of % area occupied of GFAP immunostaining normalized to control at **(B)** 1 MPI, **(C)** 3 MPI, and **(D)** 6 MPI (*n* = 11-12 animals/group). \**P*<0.05, \*\**P*<0.01, \*\*\**P*<0.001, or \*\*\*\**P*<0.0001 by two-way ANOVA with Bonnferoni’s multiple comparisons test, as indicated.

**Figure S6: Fibril-injected mice do not exhibit motor deficits or changes in anxiety-related behavior. A)** Performance in the elevated plus-maze as measured by % time in the open arms and by total number of entries into the open arms. **B)** Scoring of nest-building activity by measurement of unused nestlet material. **C)** Open-field performance measured by total distance travelled and by % time spent in the center of the apparatus. **D)** Marble-burying activity depicted by % marbles buried during assay. All assays measured at 6 MPI (*n* = 10-11 animals/group). Non-significant by unpaired Student’s *t*- test.

**Figure S7: Inter-trial performance in the Barnes maze at 1, 3, and 6 MPI. A-C)** Barnes maze performance as measured by latency to reach target hole across sequential trials at **(A)** 1 MPI, **(B)** 3 MPI, and **(C)** 6 MPI (*n* = 10-11 animals/group). \*\**P*<0.01 or \*\*\**P*<0.001 by two-way ANOVA with Bonnferoni’s multiple comparisons test, as indicated.

## Notes

### Competing Interest Statement

The authors have declared no competing interest.

