## Supplementary figures and images for "Hippocampal subfield vulnerability to α-synuclein pathology precedes neurodegeneration and cognitive dysfunction"

### Supplemental Figure 1

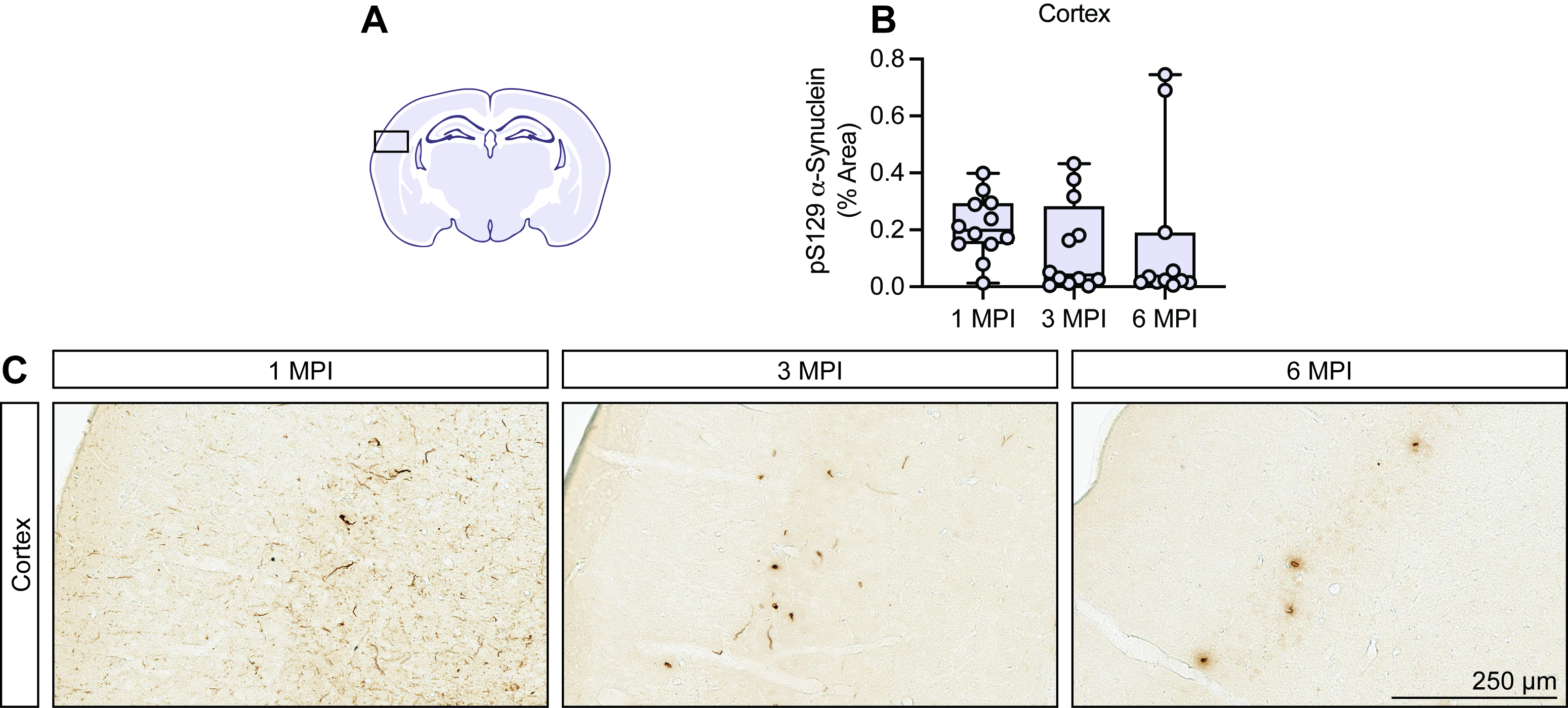

### Supplemental Figure 2

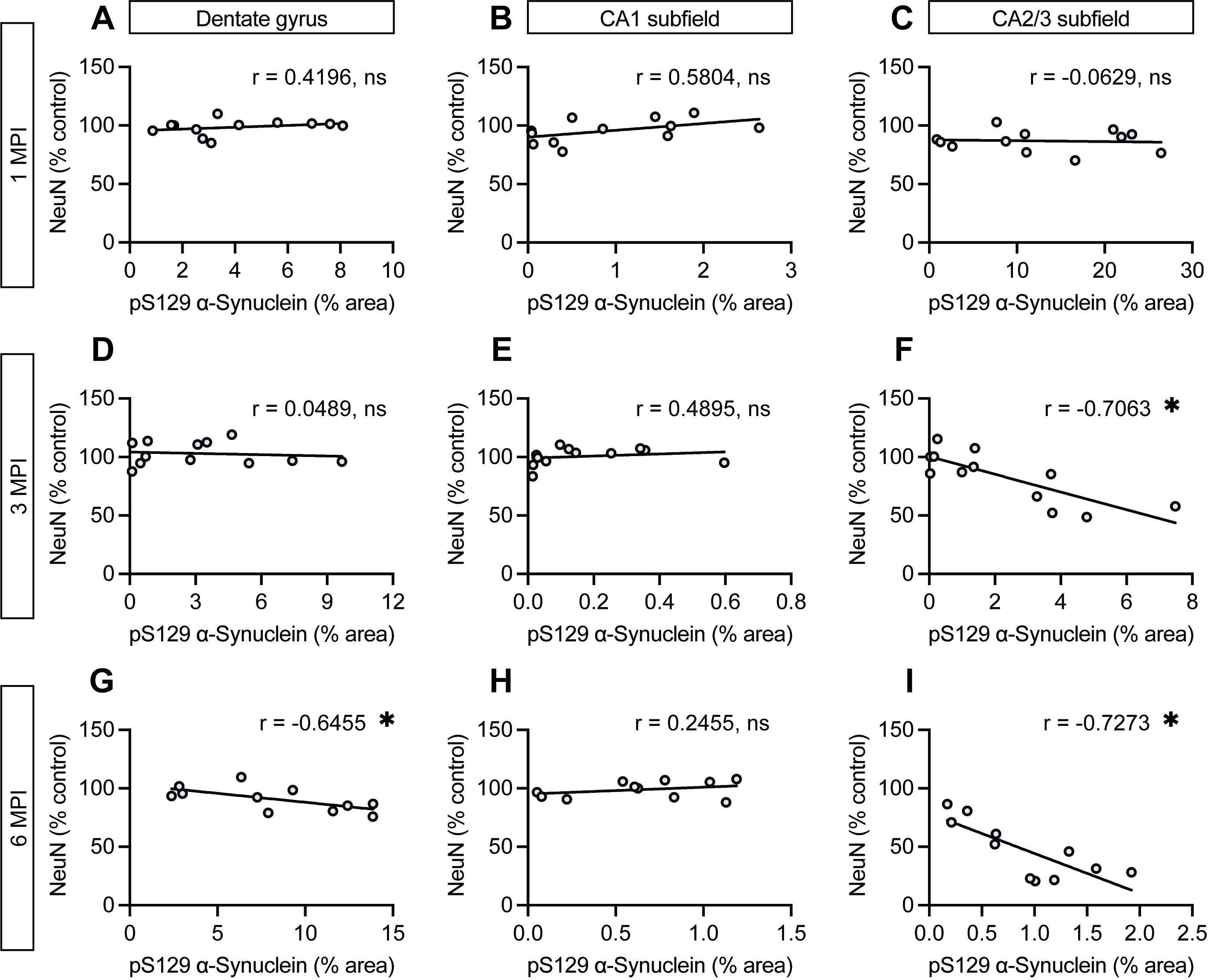

### Supplemental Figure 3

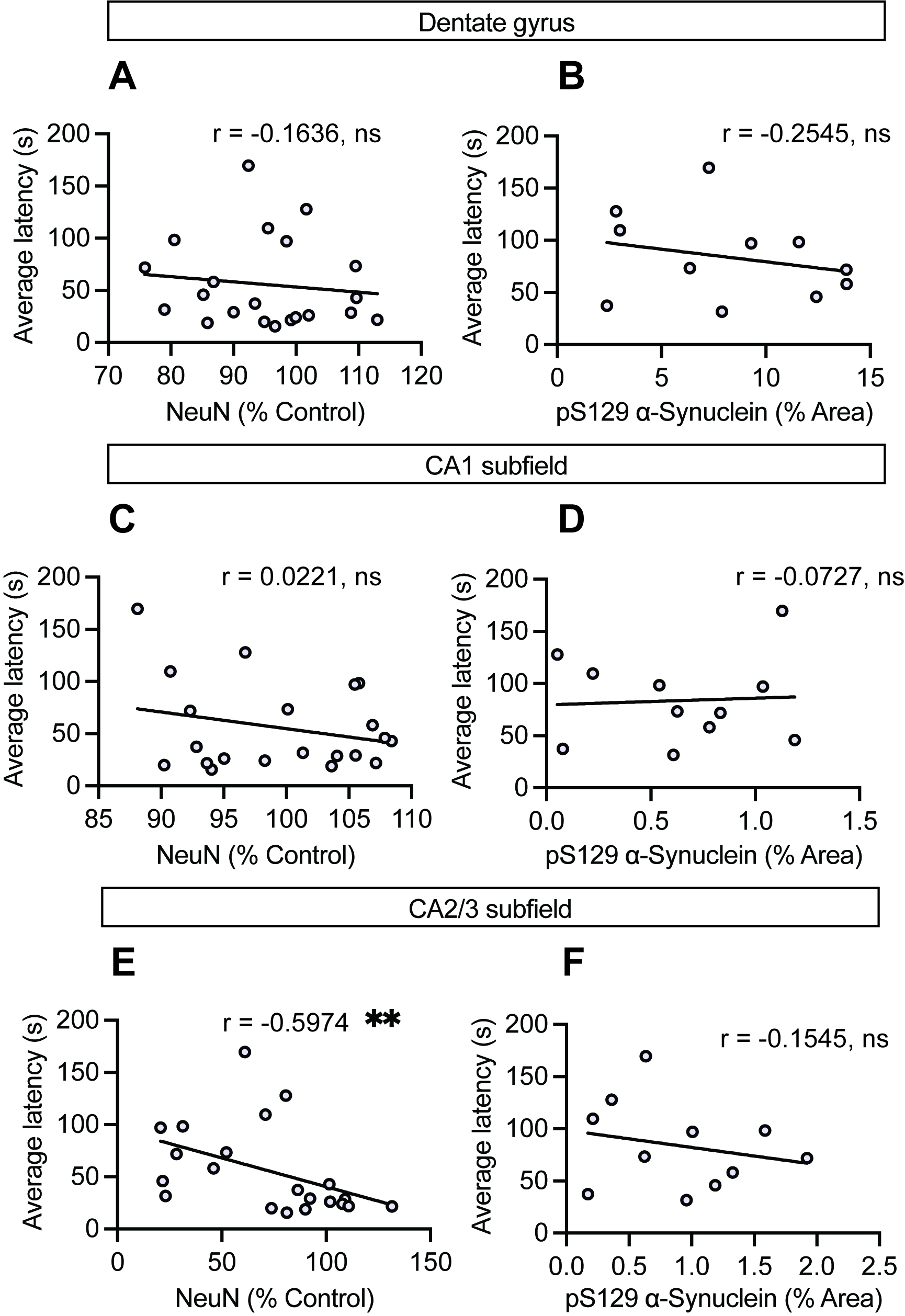

### Supplemental Figure 4

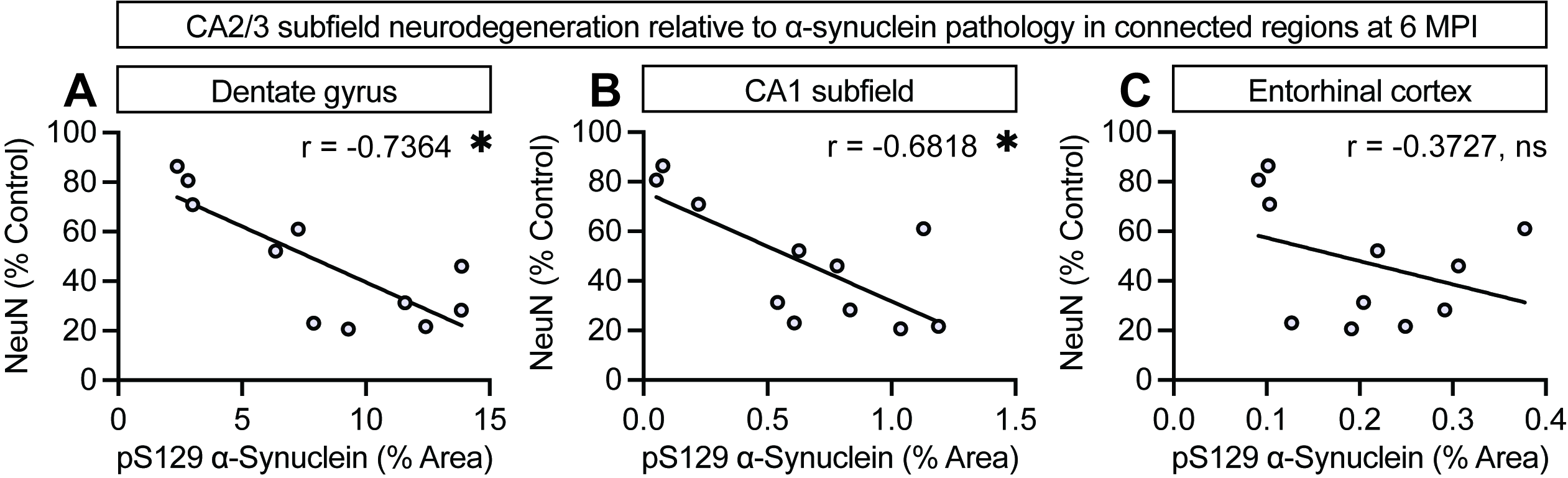

### Supplemental Figure 5

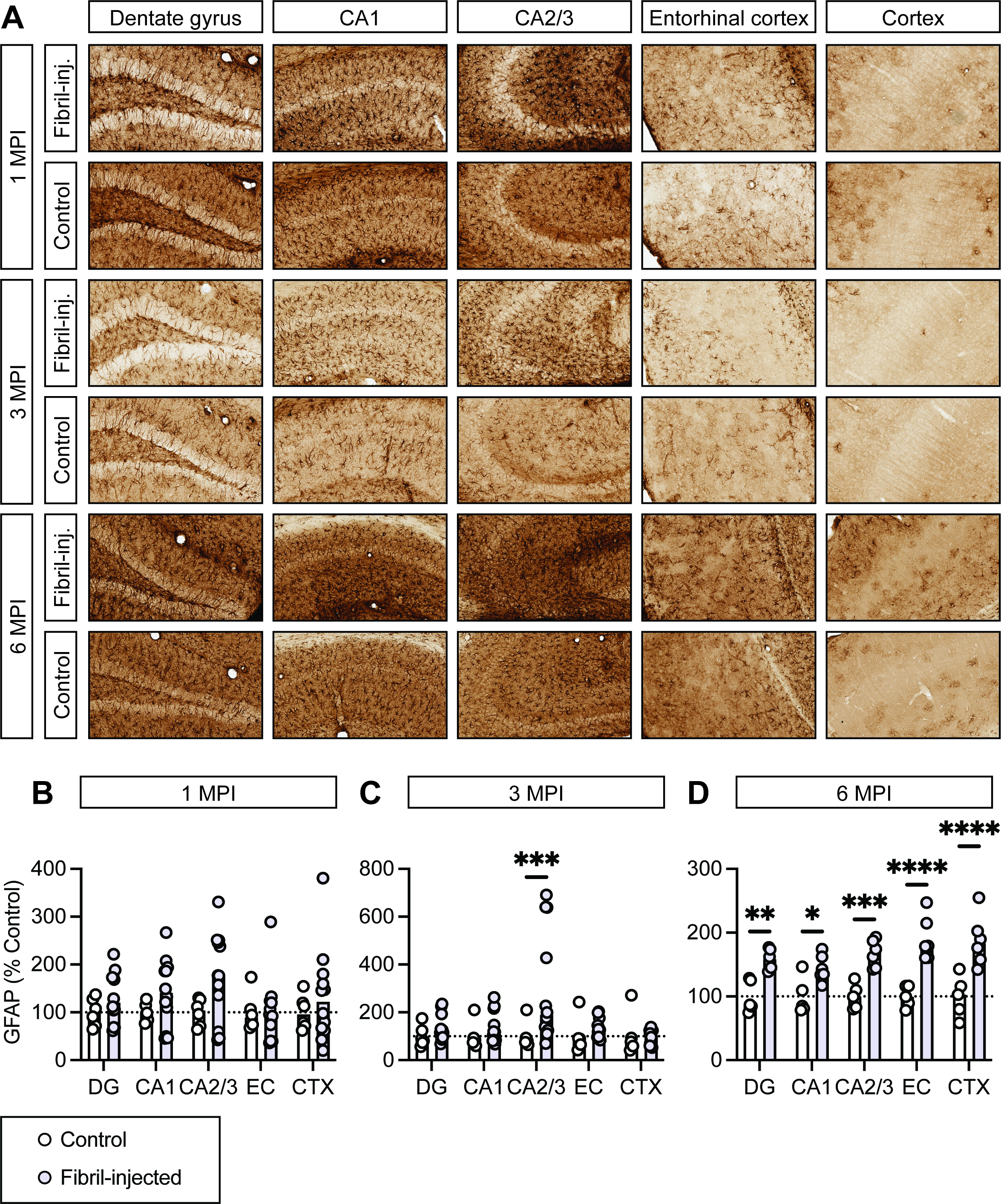

### Supplemental Figure 6

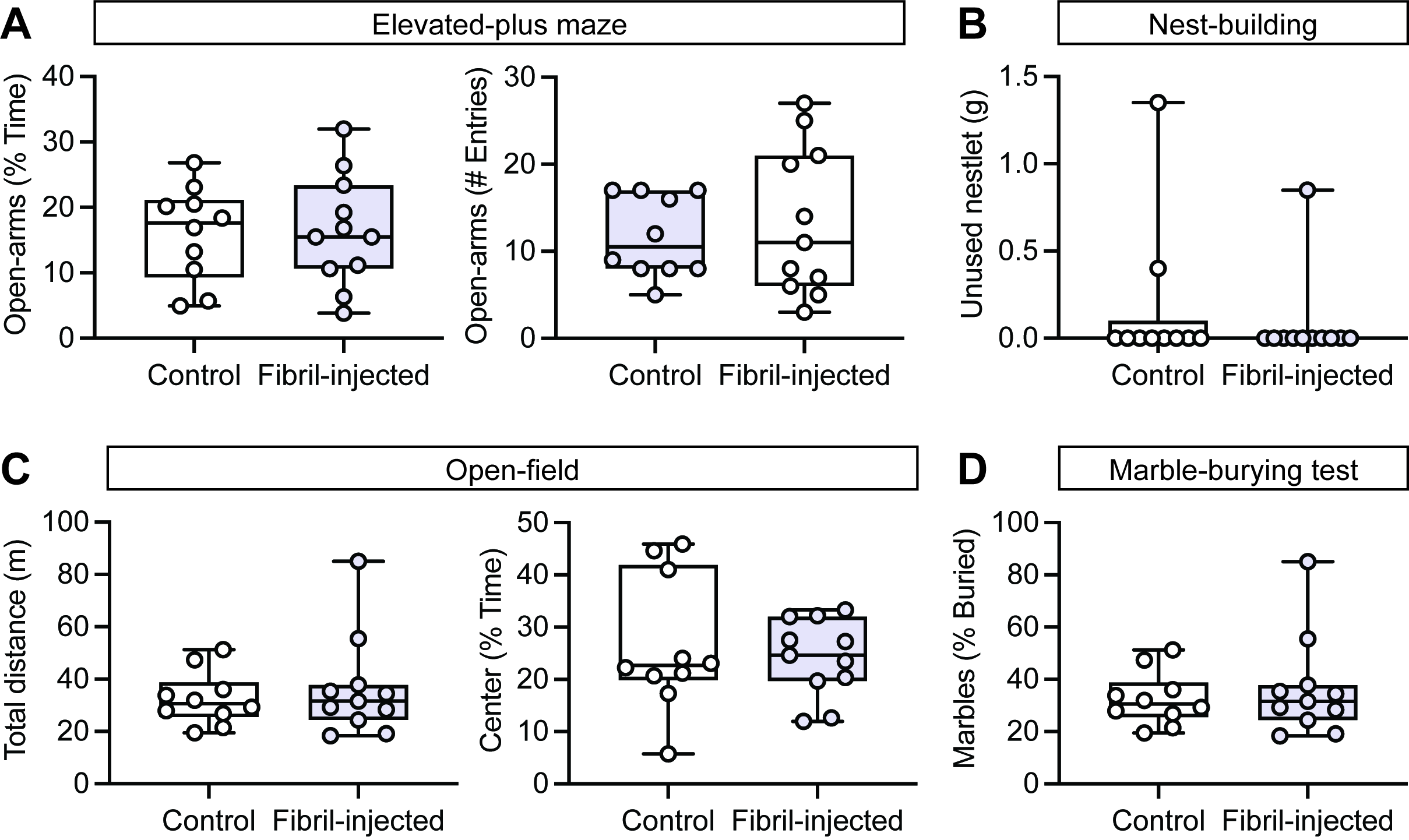

### Supplemental Figure 7

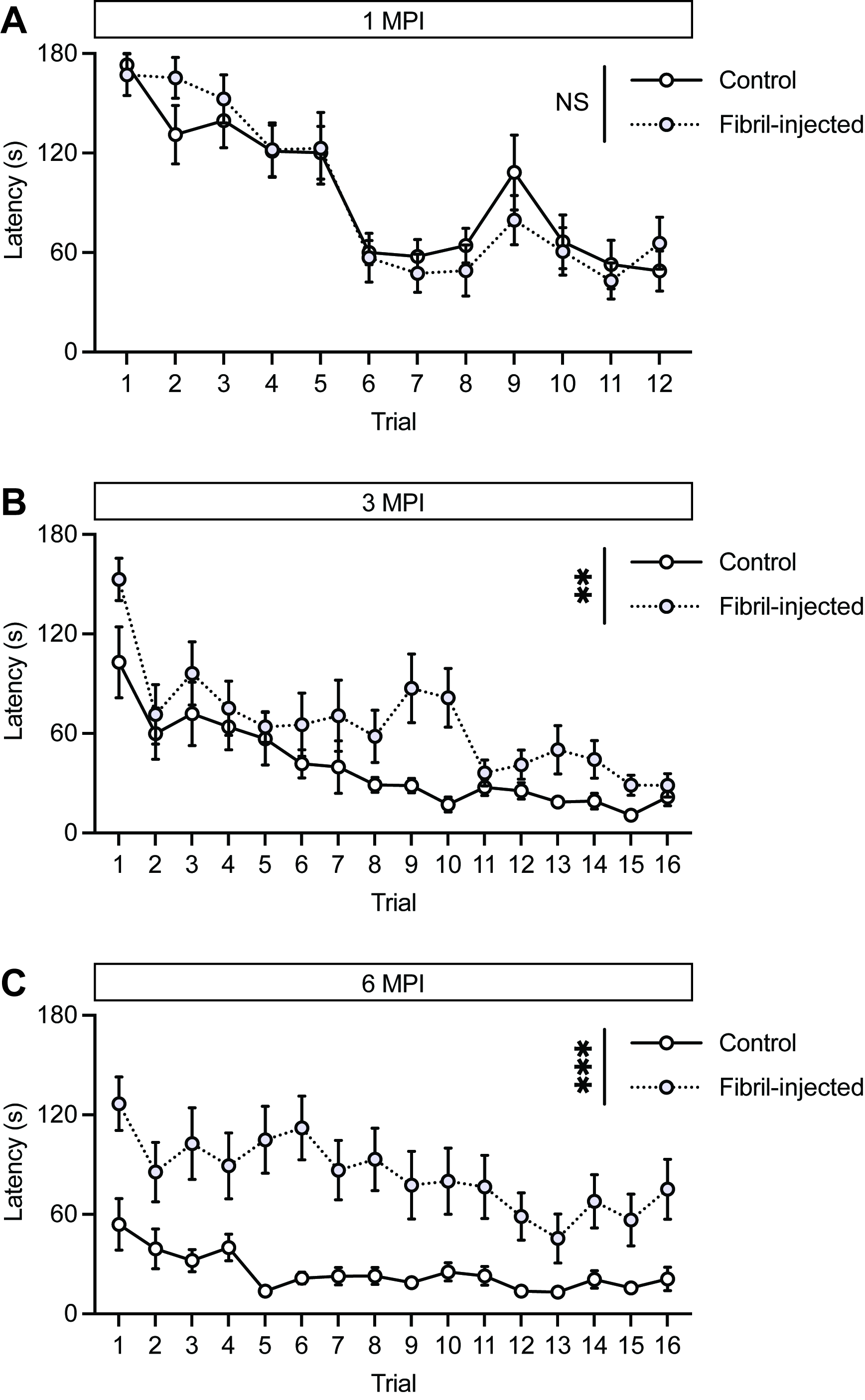
